# Flexibility and hydration of the Q_*o*_ site determine multiple pathways for proton transfer in cytochrome *bc*_1_

**DOI:** 10.1101/2024.08.22.609217

**Authors:** Sofia R. G. Camilo, Guilherme M. Arantes

## Abstract

The detailed catalytic activity of cytochrome *bc*_1_ (or respiratory complex III) and the molecular mechanism of the Q cycle remain elusive. At the Q_*o*_ site, the cycle begins with oxidation of the coenzyme-Q substrate (quinol form) in a bifurcated two-electron transfer to the iron-sulfur (FeS) cluster and the heme *b*_*L*_ center. The uptake of the two protons released during quinol oxidation is less understood, with one proton likely delivered to the histidine side chain attached to the FeS cluster. Here, we present extensive molecular dynamics simulations with enhanced sampling of side-chain torsions at the Q_*o*_ site and analyze available sequences and structures of several *bc*_1_ homologues to probe the interactions of quinol with potential proton acceptors and identify viable pathways for proton transfer. Our findings reveal that side chains at the Q_*o*_ site are highly flexible and can adopt multiple conformations. Consequently, the quinol head is also flexible, adopting three distinct binding modes. Two of these modes are proximal to the heme *b*_*L*_ and represent reactive conformations capable of electron and proton transfer, while the third, more distal mode likely represents a pre-reactive state, consistent with recent cryo-EM structures of *bc*_1_ with bound coenzyme-Q. The Q_*o*_ site is highly hydrated, with several water molecules bridging interactions between the quinol head and the conserved side chains Tyr147, Glu295, and Tyr297 in cytochrome *b* (numbering according to *R. sphaeroides*), facilitating proton transfer. A hydrogen bond network and at least five distinct proton wires are established and possibly transport protons via a Grotthuss mechanism. Asp287 and propionate-A of heme *b*_*L*_ in cytochrome *b* are in direct contact with external water and are proposed as the final proton acceptors. The intervening water molecules in these proton wires exhibit low mobility, and some have been resolved in recent experimental structures. These results help to elucidate the intricate molecular mechanism of the Q-cycle and pave the way to a detailed understanding of chemical proton transport in several bioenergetic enzymes that catalyze coenzyme-Q redox reactions.

## Introduction

Cytochrome *bc*_1_ and its homologue cytochrome *b*_6_*f* are essential proteins respectively for cellular respiration and photosynthesis. As part of electron transport chains, these enzymes balance the redox state of coenzyme-Q (quinol/quinone forms, here abbreviated by Q) in the membrane pool and contribute to the transmembrane electrochemical potential via the Q-cycle reaction.^1–4^ Cytochrome *bc*_1_ is a major producer of reactive oxygen species^5^ and is involved in various metabolic disfunctions.^6^ Inhibitors of the *bc*_1_ activity have wide biomedical and biotechnological relevance for controlling pathogens, such as in treatment of pneumonia and malaria^7^ and as commercial fungicides used in agriculture. ^8^

The structure of cytochrome *bc*_1_ is a dimer (Figure 1) and each monomer has three essential subunits: cytochrome *b* (cyt *b*), cytochrome *c*_1_ and the Rieske protein, complemented by (up to 8) organism-specific supernumerary units. Each monomer has two distinct active sites, Q_*o*_ and Q_*i*_, where two-electron oxidation and reduction of Q substrates occur. Two *b*-type hemes in cyt *b*, hemes *b*_*L*_ and *b*_*H*_ labeled after their relative *L*ow and *H*igh redox potentials, mediate electron transfer from the Q_*o*_ site to the Q_*i*_ site. A *c*-type heme in cytochrome *c*_1_ mediates electron transfer from a [2Fe-2S] cluster, also part of the Q_*o*_ site, to the soluble cytochrome *c*. This FeS cluster is bound to the Rieske protein by two histidine residues (His131 and His152 in *R. sphaeroides* numbering) and two cysteins (Cys129 and Cys149).^3,10^

**Figure 1:**
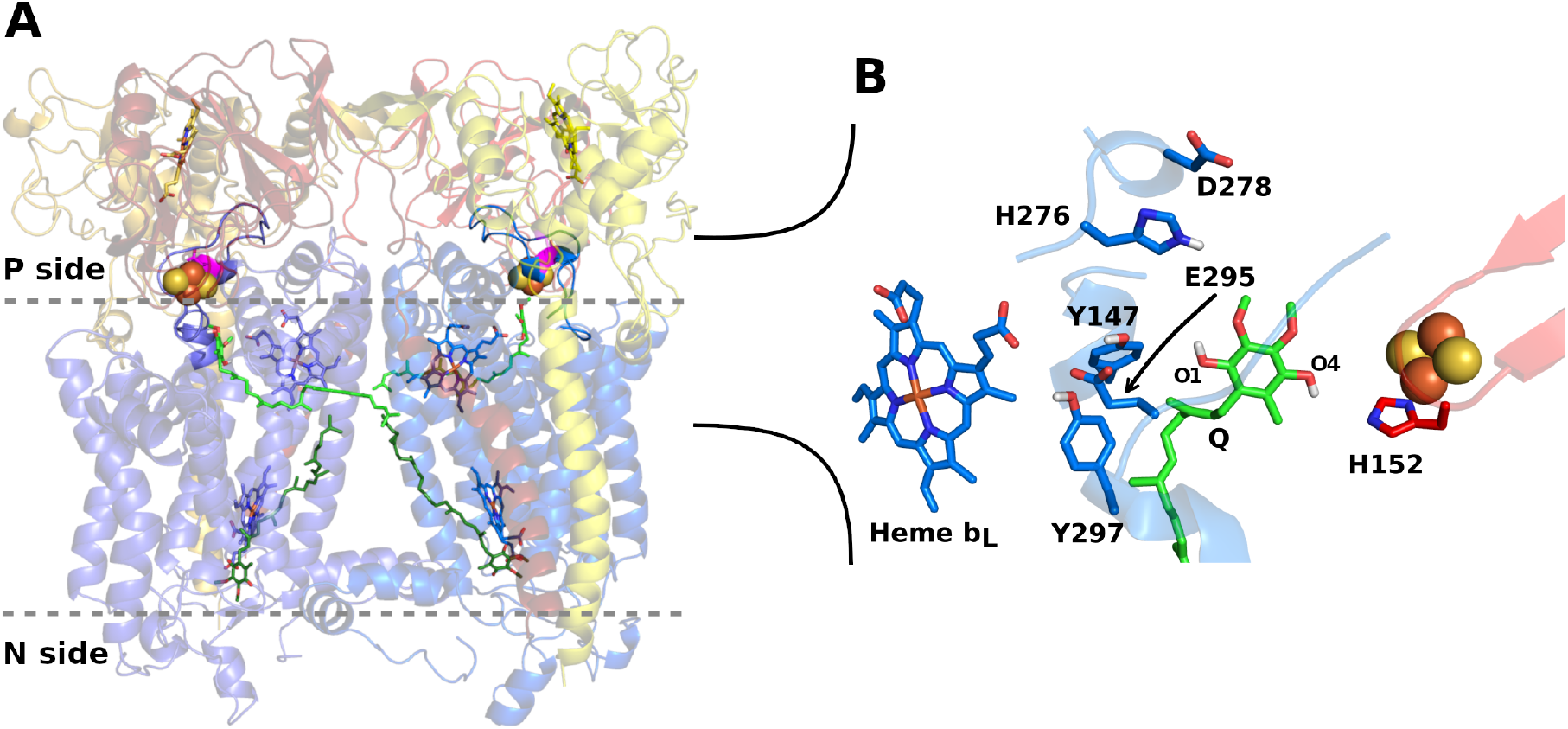
Structure of the cytochrome *bc*_1_ dimer complex of *R. sphaeroides* (PDB ID 2qjp).^9^ **A** shows essential catalytic units in the *bc*_1_ dimer in cartoon with cytochrome *b* (cyt *b*) in blue, cytochrome *c*_1_ in yellow, and Rieske protein in red. Membrane interface is in gray dashes with Q molecules in green sticks and modeled in quinol form bound to the Q_*o*_ site (membrane P side) and quinone form bound to the Q_*i*_ site (N side). FeS clusters are in orange and yellow spheres, hemes *b*_*L*_ (near site Q_*o*_) and *b*_*H*_ (near site Q_*i*_) in blue sticks, hemes *c* in yellow and cyt *b* D278 in magenta. **B** gives a close view in a rotated angle of the Q_*o*_ site with labels in residues and in phenolic oxygens of the Q substrate, which was manually modeled by replacing the inhibitor stigmatellin.

Despite its importance, the molecular mechanism of the Q-cycle catalyzed by cytochrome *bc*_1_ is still not completely elucidated.^4,11^ In the Q_*o*_ site, the two-electron oxidation of the Q substrate in quinol form is a bifurcated process,^12^ with one electron transferred to the FeS cluster with high redox potential and another electron transferred to heme *b*_*L*_. Simultaneously, two chemical protons are released from Q (resulting in the quinone form) and translocated to the membrane P side (Fig. 1). An important open question about the Q-cycle concerns the identity and the structure of protein groups in the Q_*o*_ site involved in uptake and transport of these chemical protons.

Experimental structures with atomic resolution containing a Q molecule bound to the complete Q_*o*_ site were observed only recently by cryo-EM microscopy.^13–19^ The Q substrate occupies a position similar to that previously determined for the inhibitor stigmatellin (Fig. 1B).^4,9^ The phenolic oxygen O4 in the Q-head (Q_*O*4_) and the side chain of H152 in the Rieske protein clearly interact and hydrogen-bond (H-bond) in some models.^15–18^ These observations confirm previous proposals about the essential role of H152 in substrate binding ^4^ and as the acceptor of one chemical proton.^2,20^ Direct titration of the H152 side chain in NMR experiments under different redox conditions,^21^ and various computer simulations of molecular dynamics (MD) in the Q_*o*_ site^22–26^ and of the proton-coupled electron transfer reaction between quinol and H152^26,27^ also support that this residue is one of the proton acceptors.

The binding interactions and groups involved in proton transfer from the other Q phenolic oxygen (O1) in the Q_*o*_ site are not clearly established. Single-point mutations in Y147^28^ and E295^4,29^ in cyt *b* (Fig. 1B) suggested these residues are important for Q binding and proton uptake. MD simulations have shown that Q_*O*1_ in quinol form may H-bond with side chains of Y147 and E295,^23^ and with water molecules.^22^ However, cryo-EM structures do not confirm direct interactions with these residues, which are distant by at least 0.8-1.0 nm from Q_*O*1_ in all Q bound models.^15–18^ Based on recent mutational studies, His276 and Asp278 in cyt *b* were also proposed as possible acceptors of this secondary chemical proton.^30^

Proton transport within proteins occurs through proton-wires, which are series of connected molecules that transfer an excess H^+^ from the initial donor to the bulk solvent. Water molecules and protonable groups, such as acidic side chains, are the most common components.^31,32^ A notable example is proton transport in cytochrome *c* oxidase (CcO or respiratory complex IV), where proton-wires composed of various side chains and propionate groups of redox-active heme centers have been proposed.^33–35^ Proton conduction is achieved through the Grotthuss mechanism, which involves the breaking and forming of covalent hydrogen bonds and the re-orientation of participating groups. Therefore, the composition, structure, and conformational flexibility of the participants in a proton-wire, along with their electrostatic interactions with the surrounding environment, are crucial to wire stability and transport efficiency. These properties have often been studied using molecular simulations.^36^

Here, we seek the molecular groups involved in the transport of chemical protons released by the quinol substrate in the Q_*o*_ site of cytochrome *bc*_1_. We first analyze multiple sequences of cytochrome *b* to find conserved residues spatially near the Q_*o*_ site that could act as proton acceptors. Considering the residues previously mentioned, we identify that cyt *b* Y147, E295 and Y297 are highly conserved, but H276 and D278 are not. To probe the conformational landscape and detailed interactions among these residues and the quinol substrate, we conducted molecular dynamics simulations one order of magnitude longer than previously reported^22,23,25^ employing a force-field calibrated for Q interactions that provides superior performance.^37,38^ We also employed metadynamics^39^ simulations to increase sampling of Y147, E295 and Y297 side chain torsions. These methodological enhancements led to high quality models of the molecular flexibility of side chains and the Q-head, and of internal hydration describing a network of H-bonds in the Q_*o*_ site, in agreement with the collection of experimental structures available. We identify Y147 as the initial proton acceptor from Q_*O*1_ and propose, for the first time, that the heme *b*_*L*_ propionate-A (PRA_*bL*_) is the final acceptor of the secondary proton, before releasing it to bulk water. We describe molecular wires^32^ that may transport the chemical protons released from oxidized Q via a Grotthuss mechanism^31^ and conclude that residues used by cytochrome *bc*_1_ to uptake protons may be used similarly in other respiratory enzymes that catalyze Q redox reactions.

## Methods

### Multiple sequence alignment

Starting from the protein sequence of *Rhodobacter sphaeroides* cytochrome *b* (Uniprot Q02761), a multiple sequence alignment from the region surrounding the Q_*o*_ site (residues 130 to 310) was performed using the MPI Bioinformatics Toolkit ^40^ against the Uniref90 30 jun database. This was used to determine consensus residues that could participate in catalysis by the Q_*o*_ site. All parameters were kept as default except `it max target hits` which was set to 10000. Sequences containing more than 25% gaps were removed. WebLogo was used to generate the sequence conservation plot in Fig. 2.^41^

**Figure 2:**
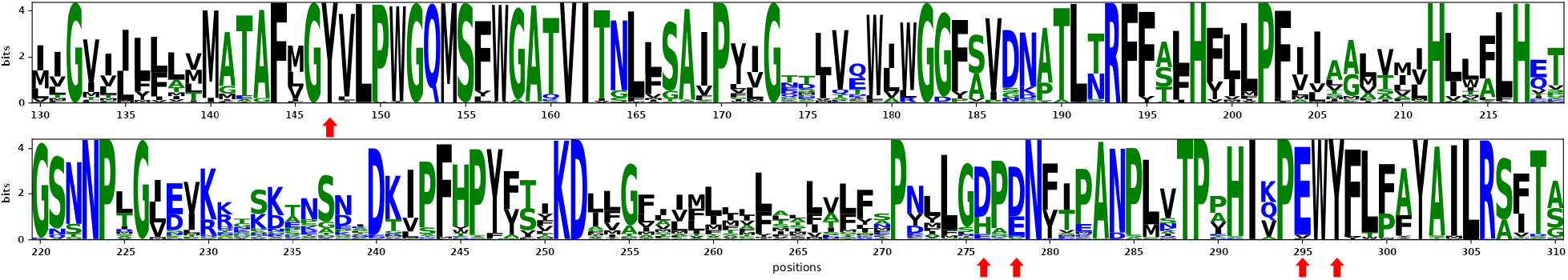
Residue conservation for cytochrome *b* shown as a WebLogo.^41^ Red arrows point to residue positions studied in the following sections (Y147, E295, Y297, H276 and D278.).

### Analysis of experimental structures

The conformational distribution of side chains in the Q_*o*_ site was analyzed by collecting 52 entries deposited in the Protein Data Bank (PDB), comprising 47 structures from cytochrome *bc*_1_ (35 from *bc*_1_ alone, 7 from supercomplex SC III-IV and 5 from SC I-III) and 5 from cytochrome *b*_6_*f* (homologous to *bc*_1_). A total of 30 models were obtained through X-ray crystallography and 22 were obtained through cryo-EM; 17 PDB entries had structural waters built into the model, and 13 entries had a Q substrate bound in the Q_*o*_ site. Only one *bc*_1_ monomer was present in some of the structures, so we analyzed a total of 79 models of the Q_*o*_ site. More details of all PDB entries and their references are listed in the Supporting Information (SI).

### Set-up of molecular models

A model of the cytochrome *bc*_1_ protein complex was constructed based on the X-ray crystal structure of *R. sphaeroides* (PDB 2qjp^9^). This model lacks subunit IV, which was recently revealed (PDB 8asi^15^) to be placed at the opposite side of the *bc*_1_ complex, more than 30Å away from the Q_*o*_ site. Thus, the lack of subunit IV in our model should not interfere with any of our simulation results and conclusions. It should also be noted that *Rhodobacter capsulatus* contains a fully active cytochrome *bc*_1_ which monomer is formed only by the three essential subunits.^4^ Inhibitors, crystallographic water, and detergent molecules were removed, while six tetra-linoleoyl cardiolipins were added, following their positions from a superimposed yeast model (PDB 1kb9^42^). Ubiquinone-6 (Q_6_, with 6 isoprenoid units) was modeled in both Q_*i*_ sites of the dimer. The oxidized form of Q_6_ was used, and its positions were adjusted through manual docking in PyMOL,^43^ with the Q-head replacing the antimycin inhibitor and isoprenoid units arranged in a U-shaped conformation. In both Q_*o*_ sites, Q_6_ was modeled in the reduced quinol (QH_2_) form, N_*ϵ*_ in H152 in Rieske protein was deprotonated, and the [2Fe-2S] cluster was oxidized. The Q molecule was manually placed, and its head replaced stigmatellin, with isoprenoid units arranged in an extended conformation. The protonation states of side chains were adjusted to positive charge in K and R residues, negative in D and E residues, and all other residues were treated as neutral, except for Asp373, exposed to the membrane in subunits cyt *b*, which was protonated. His tautomers and missing side chain atoms were assigned using WhatIf.^44^ H131 in Rieske protein, which also binds the FeS cluster, was modeled with a protonated N_*ϵ*_.^21^ The protein complex was embedded in a solvated POPC (1-palmitoyl-2-oleoylsn-glycero-3-phosphocholine) membrane with 512 lipid molecules, 39102 water molecules, 214 Na^+^ and 158 Cl^−^ ions. These ions maintain a neutral total system charge and a salt concentration of approximately 0.1 M. The complete solvated model comprised 215264 atoms.

### Classical molecular dynamics simulations

Interactions between proteins, lipids, and ions were described using the all-atom CHARMM36m force-field,^45–47^ while standard TIP3P^48^ was used to represent water molecules. Q was described using our calibrated force-field,^37,38^ and FeS clusters were represented using the Kim^49^ parameters. Oxidized heme groups were described using parameters by Luthey-Schulten *et al*.^50^ After initial geometry optimization with a conjugated-gradient minimizer, four molecular dynamics (MD) simulations of 50 ns each were performed to relax and equilibrate the complete solvated model. Harmonic restraints were applied to tether protein heavy atoms to their initial positions in the first simulation, and these restraints were successively diminished in subsequent runs, ultimately reaching zero in the final simulation. MD simulations were carried out with constant temperature (310 K) and pressure (1 atm), and a time step of 2 fs. Long-range electrostatics were treated with the Particle Mesh Ewald method.^51^ Visualization and figure plotting were performed using PyMOL^43^ and Matplotlib.^52^ A productive canonical MD trajectory was obtained with a total time of 550 ns using GROMACS version 2016.3^53^

Starting from a snapshot at 400ns of this trajectory, two additional simulations lasting 795 ns each were produced with metadynamics to enhance sampling of flexible residues in the Q_*o*_ site. In one simulation, well-tempered metadynamics^39^ was activated in the torsion of dihedrals *χ*_1_ of Y147 and Y297, and *χ*_2_ of E295 of cytochrome *b* in one monomer (chain A). The second simulation had metadynamics activated in the same dihedrals but for the other monomer (chain D). Gaussians were deposited every 1000 time steps (2 ps), at initial height of 0.6 kJ mol^−1^, widths of 0.4 and a bias factor of 15. Metadynamics was run with GROMACS version 2020.2 and the PLUMED plugin version 2.6.1.^54^

Two types of analysis were performed in the simulations. Free energy analysis is preferred because it accounts for population differences and for statistical significance of simulated processes and interactions.^55^ However, meaningful free energy profiles require extensive sampling and thus, were only used here for torsions and pair distances directly involving residues Y147, E295 and Y297. Conformational sampling for these side chains was enhanced by metadynamics, so it is expected that their torsion profiles will be the most precise (as confirmed by smaller standard errors observed in Fig. 3A-C). Sampling of pair distances was not directly enhanced so free energy profiles for distances will have lower precision. In all free energy profiles, the metadynamics bias was removed by re-weighting the distributions. Interactions involving other groups and water solvation were analyzed by trajectories of atom-pair distances and of water bridge contacts, which were considered to be formed when both distances from the bridge water oxygen to the H-bond donor and acceptor are simultaneously smaller than 0.35 nm. The six trajectories shown for each property analyzed in the Results section represent local sampling starting from different initial conformations and thus, will naturally show different histogram distributions (Count) due to the finite and localized sampling. The dispersion between these distributions is expected to be higher and the analysis of the respective property should be only qualitative.

**Figure 3:**
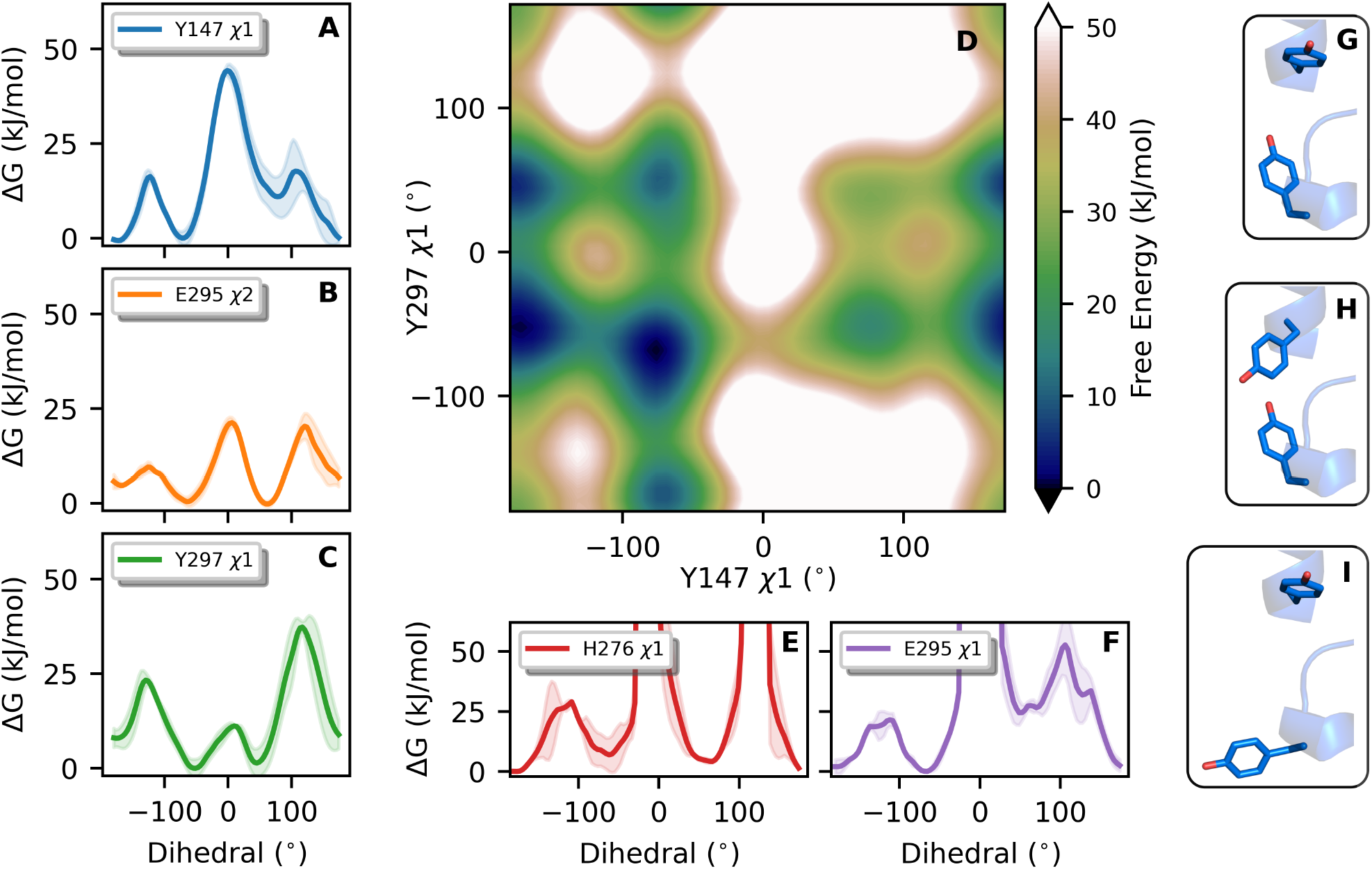
Conformational landscape of residues in the Q_*o*_ site depicted as free energy surfaces (ΔG). Side-chain torsions in cyt *b* residues indicated in the legend are shown in **A**-**F**, with a 2D-profile shown in **D** for *χ*1 in both Y147 and Y297. 1D-profiles show the average of two independent metadynamics simulations, with colored shadows representing the standard error. Torsions of dihedrals *χ*1 in Y147 and Y297, and *χ*2 in E295 (**A**-**C**) were boosted in the metadynamics, while torsions of *χ*1 in H276 and in E295 were not. Thus, discontinuities in **E** and **F** represent undersampled regions. Illustrative structures for (Y147, Y297) side chains are shown respectively for conformers (*t, g*) in **G**, (−*g, g*) in **H** and (*t*, −*g*) in **I**.

## Results & Discussion

### Y147, E295, and Y297 in cyt *b* are highly conserved while H276 and D278 are not

Protein residues with high conservation along the evolutionary process are expected to have important structural and catalytic roles. We employed a multiple sequence alignment (MSA) to explore residue conservation in cytochrome *b* (9,993 sequences with 1.36e-123 *<* e-value *<* 2.00e-51 from the *R. sphaeroides* sequence) and found that most of the residues in the Q_*o*_ site are highly conserved, as shown graphically in Fig. 2. Focusing in side chains that may change protonation state and participate in proton transport near the Q binding site, residues Y147, E295 and Y297 (here named as YEY group) show 99.9%, 94.2% and 98.9% conservation, respectively, raising to 100%, 99.7% and 100% among 329 sequences curated in the Swiss-Prot database as active enzymatic subunits. ^56^ For comparison, residues Pro294 and Trp296, in the often studied and conserved PEWY motif,^57–60^ are conserved in 99.3% and 99.0% sequences in the full set. No double mutations among the YEY residues were observed, which could indicate a co-evolutionary dependence.

H276 and D278 have much lower conservation, 23.4% and 68.3% in the full set, and 10.2% and 80.1% for the sequences in Swiss-Prot, suggesting that these are not essential residues for function. However, these positions are often substituted by acidic residues (H276D and D278E) which may also transport protons. Thus, a more detailed analysis is necessary to determine the role of H276 and D278 in the Q_*o*_ site.

Although we focused on high similarity (low e-values were used to determine our MSA), our conclusions on residue conservation are comparable to those obtained by Hunte *et al*., who used a more diverse and less similar sequence set focused on the PEWY motif.^60^ Their study suggests parallel evolution of the Q_*o*_ site at position 295, occupied by glutamate when the substrate is ubiquinone and aspartate when the substrate is menaquinone, with lower redox potential. In position 297, only mutations from Tyr to Phe were observed in their sequence set. The respective organisms always possess extra cyt *b* sequences, suggested to perform different function such as thiosulfate oxidation. ^60^

### MD simulations unveil the flexibility of conserved residues and of the substrate Q-head in the Q_*o*_ site

The conformational landscape and intermolecular contacts of the Q_*o*_ site bound with the Q substrate were explored by three long molecular dynamics simulations modeling the reactant Michaelis complex of cytochrome *bc*_1_. The expected charge state before electron and proton transfers was assigned as the Q substrate in quinol form (doubly reduced and doubly protonated) with heme *b*_*L*_ and the [2Fe-2S] cluster in oxidized form. The H152 side chain was bound to the FeS cluster (via N*δ*) and deprotonated (at N*ϵ*). Two simulations of 795 ns had metadynamics activated to enhance sampling of torsion angles *χ*1 of Y147, *χ*2 of E295, and *χ*1 of Y297 in a single Q_*o*_ site. Because cytochrome *bc*_1_ is a dimer, two canonical MD trajectories of the same length (795 ns) for the other monomer not activated by metadynamics, were also produced. These add to the initial MD simulations of 550 ns, giving a total of four canonical MD trajectories of the Q_*o*_ site (2*×* 550 ns + 2*×* 795 ns). All simulations were stable and maintained the global fold and backbone configuration of protein subunits, even when metadynamics was activated (Figs. S1 and S2 in SI).

Side-chains in the YEY group and H276 populate *t* (*χ* angle ∼ 180^*?*^, defined using the quartet N-C*α*-C*β*-C*γ*), *g* (60^*?*^), and −*g* (−60^*?*^) conformers, indicating high flexibility in the Q_*o*_ site, as shown in the obtained torsion profiles (Fig. 3). For Y147 and Y297, five pairs of conformers may be populated (blue regions in Fig. 3D), with the most likely pairs being (*t*, −*g*) and (*t, g*), illustrated in Fig. 3G,I. Barriers for transitions between these populated conformers, such as from −*g* → *t* of the Y147 side chain, are between 10-25 kJ/mol. Thus, interconversions occur on the ns-time scale and should not gate proton and electron transfers. As discussed in the section “Experimental structures confirm that Q_*o*_ side chains are flexible and highly hydrated”, analysis of tens of *bc*_1_ structures deposited in the PDB confirms that the multiple conformations of the YEY group side chains obtained here are also experimentally observed.

Interactions of the bound substrate Q-head in the Q_*o*_ site were identified through distance trajectories taken from MD simulations. Fig. 4A shows the minimum distance between the quinol phenolic oxygen O_1_ and heme *b*_*L*_ is always under 1.5 nm. Thus, conformations probed in simulations are capable of fast electron transfer between the Q substrate and heme *b*_*L*_. In Fig. 4B, the distance between O_4_ and H152 N_*ϵ*_ is consistently lower than 0.6 nm, except in one metadynamics trajectory (colored green) between 250 to 680 ns, when the Q-head bends towards heme *b*_*L*_ and O_4_ moves away from H152. A direct H-bond is established between O_4_ and H152 N_*ϵ*_ when their distance is 0.35 nm, and a bridged interaction by one water molecule is established when their distance is 0.5-0.6 nm (a detailed description of local solvation and water-bridge contacts is given below in the section “Q_*o*_ site is highly hydrated and has several water-bridged connections between side chains”). A similar observation is made from Fig. 4C, where three characteristic distances between quinol Q_*O*1_ and Y147_*OH*_ are observed, corresponding to the three peaks at 0.3, 0.5, and 0.9 nm in Count distributions. A direct H-bond and a one-water bridge interaction are observed for distances of 0.3 and 0.5 nm, respectively (Fig. 5B-E).

**Figure 4:**
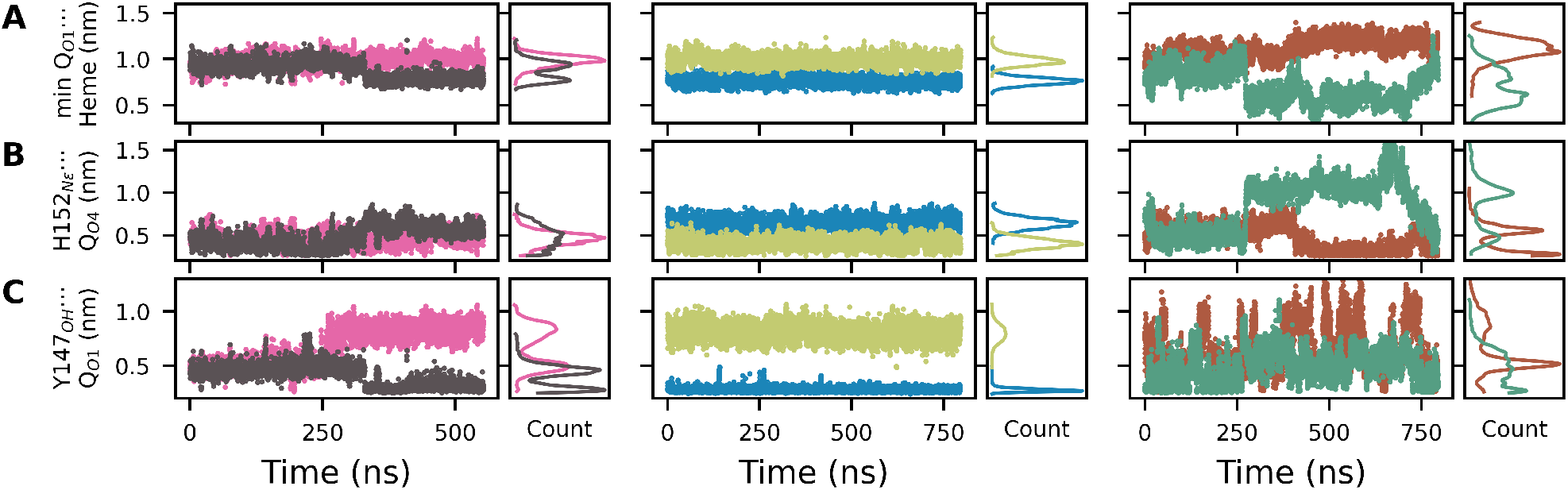
Atom-pair distances involving the substrate Q-head in the Q_*o*_ site during simulations. Pairs are given in Y-axis labels. Left and middle columns show canonical MD simulations, right column shows metadynamics simulations. Each column shows two trajectories corresponding to the two Q_*o*_ sites of the *bc*_1_ dimer. Count shows a histogram of the respective distance.

**Figure 5:**
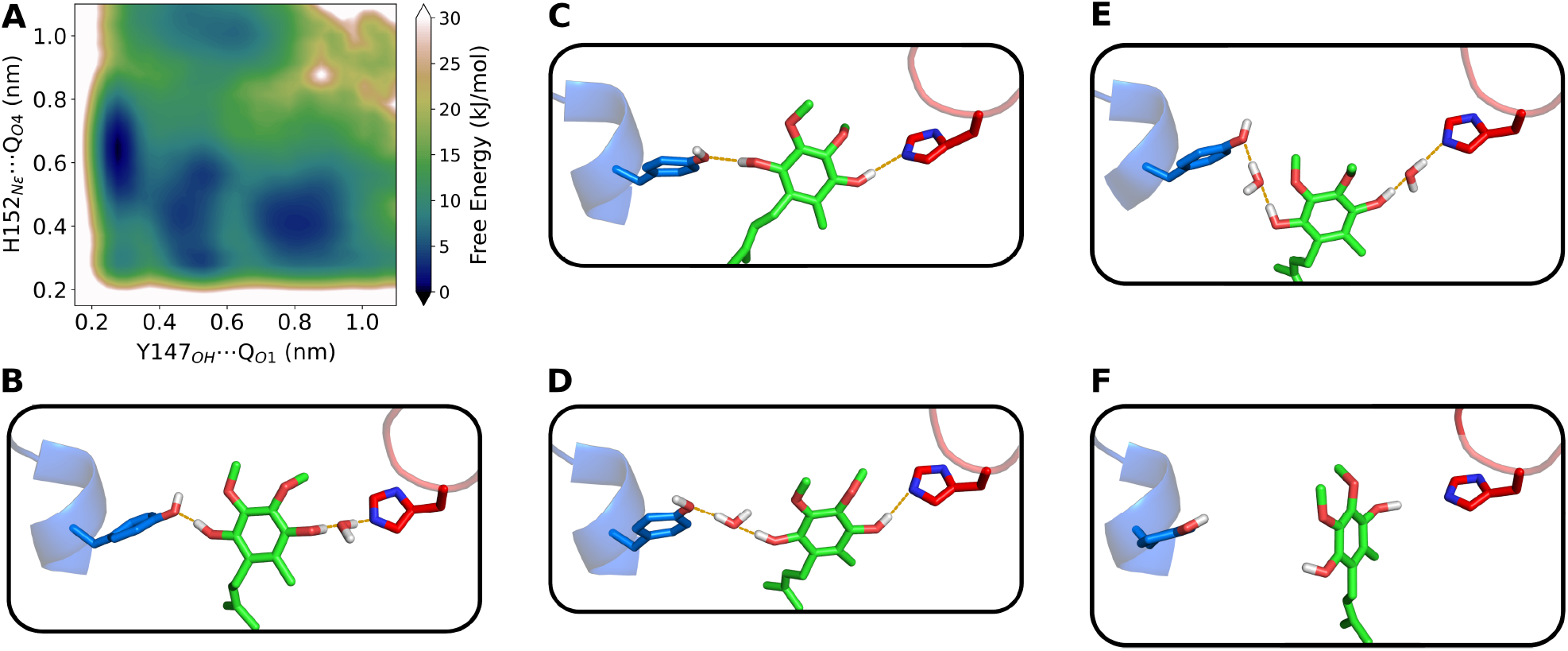
Conformational landscape of the substrate Q-head bound in the Q_*o*_ site. **A** shows the free energy surface in relation to distances between Q substrate phenolic oxygens and Q_*o*_ site residues. Representative snapshots of Y147 (blue), H152 (red) and the Q-head (green) conformations with H-bonds displayed as orange dashes are shown for distance pairs (Y147_*OH*_ *…*Q_*O*1_, H152_*Nϵ*_ *…*Q_*O*4_) in nm: **B** (0.3,0.6), **C** (0.3,0.3), **D** (0.5,0.3), **E** (0.5,0.5) and **F** (0.8,0.4). The Y147 side-chain is shown in the *t* form for all snapshots, but its −*g* conformer may also establish H-bonds similar to those in **B, D** and **E** models.

Multiple minima for the interaction of the Q-head with Y147 and H152 side chains in the Q_*o*_ site are revealed in the free energy profile shown in Fig. 5A, suggesting that the bound Q substrate is also highly flexible. This profile was obtained by combining the six trajectories of Fig. 4B-C, with an aggregate time of 4.3 *µ*s of MD simulation.

Three binding modes were identified with similar stability, corresponding to free energy minima (dark blue regions in Fig. 5A) found at Y147_*OH*_ *…*Q_*O*1_ distances of 0.3, 0.5, and 0.85 nm. The first mode (at 0.3 nm, “proximal” to heme *b*_*L*_) is the most stable and is separated from other minima by interconversion barriers of 10 to 20 kJ/mol. It is characterized by a direct H-bond between Q_*O*1_ and Y147_*OH*_ and a water-bridged interaction between Q_*O*4_ and H152_*Nϵ*_ (Fig. 5B). The second mode (at 0.5 nm, intermediate to heme *b*_*L*_) is broad and the least stable among the three minima. It is characterized by a water bridge between Q_*O*1_ and Y147_*OH*_, but the interaction of Q with H152 is either a direct H-bond (Fig. 5D) or a water bridge (Fig. 5E). These two binding modes are reactive configurations that could readily transfer phenolic protons during Q oxidation, suggesting that Y147 in cyt *b* and H152 in the Rieske protein are the initial acceptors of protons released.

The third binding mode (at 0.85 nm, “distal” to heme b_*L*_) is characterized by a weak interaction between Q_*O*4_ and H152_*Nϵ*_ and a lack of contact between Q_*O*1_ and Y147_*OH*_ (Fig. 5F), disabling proton transfer between these groups. This minimum is separated from the previous two minima by a ∼10-15 kJ/mol barrier, making this third mode relatively stable.

Remarkably, this distal mode corresponds to the Q binding mode found among all experimental *bc*_1_ structures deposited in the PDB in which a Q substrate binds to the Q_*o*_ site.^13–19^ These PDB structures, obtained recently from various organisms and with substrates of variable Q-tail length, show distances of H152_*Nϵ*_ *…*Q_*O*4_ between 0.3-0.5 nm and Y147_*OH*_ *…*Q_*O*1_ between 0.8-1.0 nm, in line with the mode in Fig. 5F. Thus, we suggest that the distal mode represents a stable pre-reactive conformation that could be observed in typical cryo-EM experiments,^13–16^ while the other two more proximal modes (Fig. 5) are in dynamic exchange and cannot be captured by cryo-EM due to the high flexibility and proposed reactivity (short lifetime during the Q-cycle) of the Q substrate.

The three binding modes are observed for both −*g* and *t* conformers of the Y147 side chain, while its *g* form often leads to slight dissociation of the Q-head from the Q_*o*_ site, disrupting the contact between Q_*O*4_ and H152. The configuration with a double H-bond found in the distance pair (0.3,0.3) nm shown in Fig. 5C has low stability and was not considered a stable binding mode. It was observed in simulations only when Y147 was in the *t* form. Thus, Y147 flexibility modulates the binding interactions of the Q-head and its contacts with H152.

The minimum in the profile centered at pair (0.6,1.0) nm corresponds to the Q-head bent towards heme *b*_*L*_, observed only in a stretch of one metadynamics trajectory (mentioned above). This region does not represent a Q binding mode and may be an intermediate for the substrate binding pathway, such as proposed for Q binding in other respiratory enzymes.^61,62^

The stability of other direct contacts involving YEY side chains was evaluated in Fig. 6, with free energy profiles estimated from the metadynamics simulations. The interaction between the O_1_ phenolic oxygen in Q and the E295 side chain (C*δ* is used as a reference since both O*ϵ* can form H-bonds) is weak (minimum *<*5 kJ/mol) and these groups are more stable when separated by ∼0.7 nm (Fig. 6A). Side-chains of Y147 and E295 are highly flexible (Fig. 3), but their inter-residue contact is stable (minimum at 0.5 nm in Fig. 6B).

**Figure 6:**
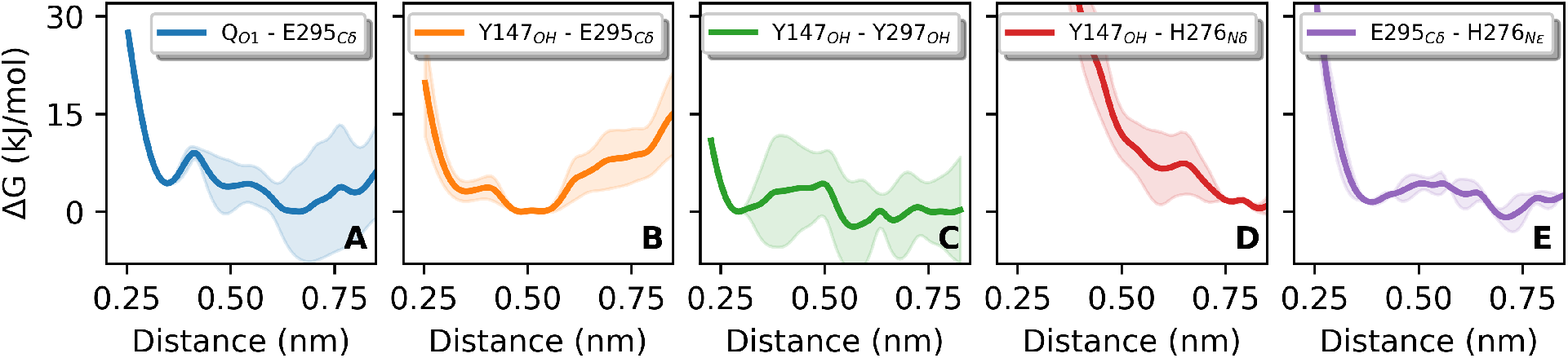
Stability of direct interactions in the Q_*o*_ site depicted as free energy (ΔG) profiles for the contacts indicated in the legend of each panel. Lines show the average of two independent metadynamics simulations and colored shadows represent the standard error.

The Y147 and Y297 side chains can also H-bond (minimum at 0.3 nm in Fig. 6C), but it is difficult to quantify their stability because this free energy profile has a large variance and is less reliable at longer pair distances. This H-bond is only formed when Y147 and Y297 residues are in the (−*g, g*) conformer (Fig. 3H). Heme *b*_*L*_ A-propionate (PRA_*bL*_) does not (or very rarely) form direct H-bonds with side chains of the YEY group (Fig. S4A-C). However, PRA_*bL*_ can interact with the YEY residues via multiple water bridges, as depicted in the next section.

The role of H276 in proton transfer from the Q substrate^30^ can be evaluated from its contacts in the Q_*o*_ site. H276 does not interact directly or via a water bridge with either the substrate Q-head (Fig. S4D and Fig. S5F) or residue Y147 (Fig. 6D and Fig. S5G). However, H276 and E295 form a weak inter-residue contact (minimum *<* 5 kJ/mol, Fig. 6E). Thus, H276 should not receive a proton directly from the Q-head or from the proposed initial acceptor Y147, and may participate in the proton transfer pathway only if receiving it from E295. The connection between H276 and D278, another proposed proton release group,^30^ can be easily established directly (Fig. S4E) of via water bridges (Fig. S5C,D).

### Q_*o*_ site is highly hydrated and has several water-bridged connections between side chains

Fig. 7A shows a snapshot of the complete and equilibrated *bc*_1_ model used in our MD simulations, with the solvated and membrane-embedded protein. It is evident that water can penetrate the Q_*o*_ site and connect the bulk to PRA_*bL*_ and to D278. This allows water to transit quickly and ensures high hydration within the Q_*o*_ site. Fig. 7B illustrates the average local hydration in MD simulations, as quantified in Fig. 7C. For instance, acidic oxygens in E295, D278, and heme PRA_*bL*_ are highly hydrated, with a median of 5 to 6 water molecules each. The extensive solvation of D278 and PRA_*bL*_, which are in contact with external water, suggests that these groups may function as proton release groups, i.e., as the last protein acceptors of the chemical proton before it is transferred to bulk water.

Polar oxygens in residues Y147, Y297, and in the quinol substrate (O_1_ and O_4_) are hydrated by only one molecule each on average (Fig. 7B,C). However, significantly higher water H-bond lifetimes (*t*_*life*_, calculated from the canonical MD simulations) reaching 4 ns for Q_*O*1_ and 0.4-0.3 ns for the other three groups are observed for these groups. For the H-bond between Q_*O*1_ and water, *t*_*life*_ is three orders of magnitude longer than simulated with the same energy model for a free and unbound quinol molecule embedded in lipid-only bilayers.^37,38^ On the other hand, *t*_*life*_ for water H-bonded to D278 and PRA_*bL*_, which have higher hydration and are in direct contact with the bulk, are 0.1 ns or lower and similar to bulk water *t*_*life*_. This suggests that the Q_*o*_ site stabilizes local hydration to facilitate the formation of proton-conducting wires, with water diffusion increasing with distance from the initial proton donor (Q substrate here).

**Figure 7:**
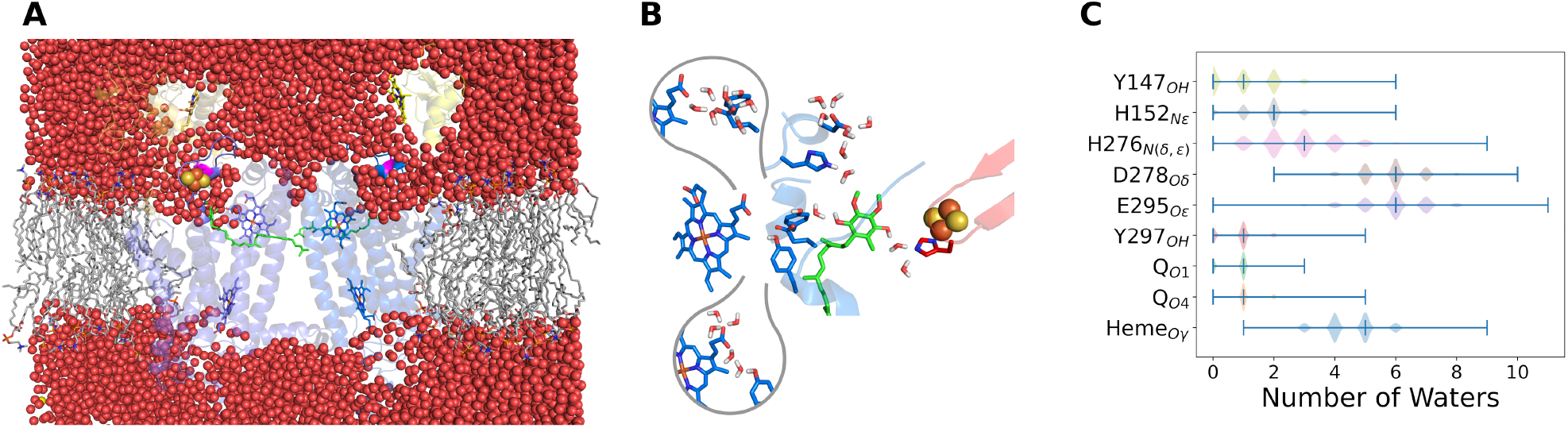
Hydration of the cytochrome *bc*_1_ dimer complex. **A** shows the global solvation of the protein model embedded in the membrane with lipids in gray sticks, in the same orientation and color code of Fig. 1A. Water molecules (oxygen in red spheres) penetrate and solvate the Q_*o*_ site, with direct contact from bulk water to heme PRA_*bL*_ (blue sticks) and to D278 (magenta). **B** shows a close view of the Q_*o*_ site with average number of bound water molecules found in simulations, in the same orientation of Fig. 1B. The two bubble insets focus on hydration of PRA_*bL*_ and YEY residues which were removed from the main panel for clarity. **C** gives the distribution of the number of water molecules under H-bond distance to centers in the Q_*o*_ site listed in the Y-axis as obtained from the MD simulations. Minimum, median and maximum numbers for each center are shown as ticks.

Several contacts between the substrate, binding residues and heme *b*_*L*_ in this highly hydrated Q_*o*_ site are mediated by water molecules in bridge. These contacts are essential to establish proton-wires that may be used in a Grotthuss mechanism of proton transfer. Figs. 8 and S5 show trajectories during MD simulations for contacts that may mediate proton transfer from the Q substrate to heme and to D278. Due to their boosted potential, metadynamics simulations (last panel column in Figs. 8 and S5) show more frequent contact exchange and a more even histogram between simulations of the two Q_*o*_ sites in the *bc*_1_ dimer, while canonical MDs (first and second panel columns in these Figures) have less exchange and more dispersion, with the dynamics of contacts depending more on their (different) initial configuration.

**Figure 8:**
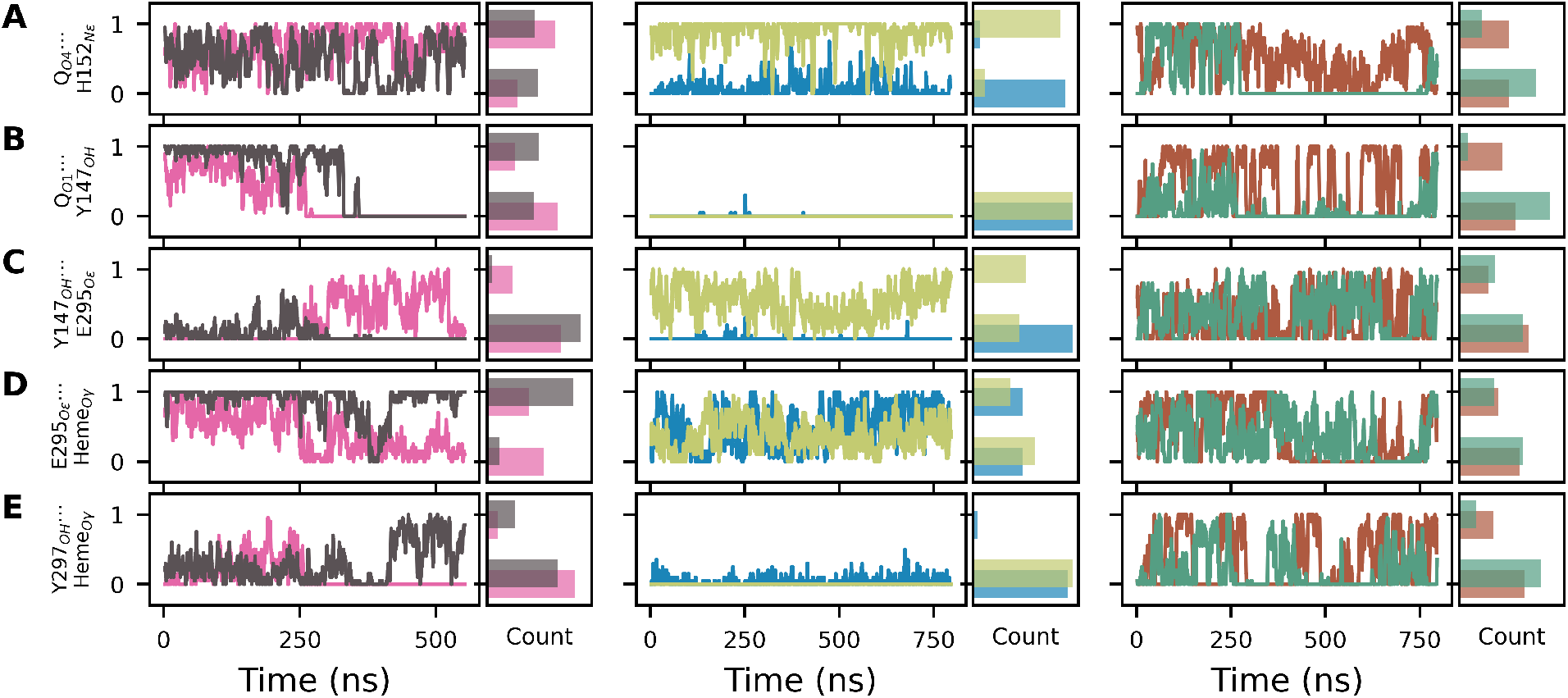
Contacts bridged by one water molecule during simulations. Panel columns and colors relate to the six MD simulations of the Q_*o*_ site as described in Fig. 4. Contacts are given in Y-axis labels and a bridge water is denoted by dots (*…*). One or zero correspond to the contact formed or not, respectively. A moving-average with a 1 ns window is plotted. Count shows a histogram of contact formation.

Bridges are frequently and briefly disrupted when the bridged water exchanges with another water molecule (fast oscillations in Fig. 8), but bridges are disrupted persistently when their respective donor-acceptor centers move away. For instance, the bridge Q_*O*4_ *…*H152_*Nϵ*_ is disrupted at ∼ 250 ns in the green trajectory of Fig. 8A, corresponding to the increase in distance between these centers in the same trajectory shown in Fig. 4B. A similar observation can be made for the bridge Q_*O*1_ *…*Y147_*OH*_ (compare Fig. 8B to Fig. 4C). Here, three characteristic distances are observed (in agreement with the minima found in the free energy profile of Fig. 5A), and the water-mediated bridge can only be established when the distance between Q_*O*1_ and Y147_*OH*_ is ∼0.5 nm. Bridged contacts involving phenolic oxygens (O_1_ and O_4_) in Q are the most persistent, with an average lifetime of 0.4 ns, while other bridge contacts involving putative proton acceptors remain for less than 0.2 ns.

Bridged contacts between Y147 and E295, and between heme PRA_*bL*_ and E295 or Y297, can often be formed (Fig. 8C-E), suggesting a combination of possible proton-wires to deliver a chemical H^+^ to heme. Interestingly, a direct (Fig. 6A) or a bridged water contact between E295 and the Q-head are not (or very rarely) established, and E295 should only participate in proton-wires that transfer a chemical H^+^ from the Q substrate via Y147 (Fig. S5E). Additional bridged contacts of heme *b*_*L*_ with Y147_*OH*_ via two sequential waters, and with Q_*O*1_ via three sequential bridged waters, may also be established (Fig. S5A,B). H276_*N*(*ϵ,δ*)_ can also connect to D278_*Oδ*_ via bridges with one or two water molecules (Fig. S5C,D).

Thus, a dynamic network of H-bonds is established in the Q_*o*_ site, as all bridged contacts shown in Fig. 8 remain formed during a non-negligible fraction of the simulated time. For instance, the least stable bridge in this figure, Y297_*OH*_ *…*Heme_*Oγ*_, is established 16% of the simulation time.

It should be noted that previous MD simulations suggested a mostly dry Q_*o*_ site that lacked an H-bond network mediated by water molecules^23,25^ in disagreement with the present results and with recent cryo-EM structures (Fig. 11). Another MD simulation included the participation of water in the Q_*o*_ site but dismissed a role for Y147 and emphasized E295 as an initial proton acceptor.^22^ We attribute these differences compared to the present results to two important methodological improvements in our study. The force field used to describe the Q substrate in previous simulations^22,23,25^ has been shown to contain errors in torsional potentials and electric dipoles^37^ and led to inaccurate partition free energies compared to experiments,^38^ possibly due to exaggerated hydrophobicity of the Q-head. Here, we employed a force field calibrated for Q interactions, which has been shown to provide superior performance^37,38^ and thus, has become the *de facto* standard to simulate Q molecular dynamics.^62–64^ We also performed simulations one order of magnitude longer (in aggregated time) and had conformational sampling enhanced by metadynamics, allowing significantly more sampling of important interactions in the Q_*o*_ site.

### Multiple proton-wires are identified and connect the Q substrate to various proton acceptors

By combining this hydration analysis with the direct contacts and side chain flexibility described in the previous section (Figs. 3 and 6), we suggest multiple proton-wires to transfer the two chemical H^+^ released after quinol oxidation in the Q_*o*_ site (Fig. 9). Three different wires connect Q_*O*1_ to heme PRA_*bL*_, one wire connects Q_*O*1_ to D278, and one wire from Q_*O*4_ to H152. Y147 is the initial acceptor from Q_*O*1_ in all proposed proton-wires, and H152 is the initial (and only) acceptor from Q_*O*4_.

**Figure 9:**
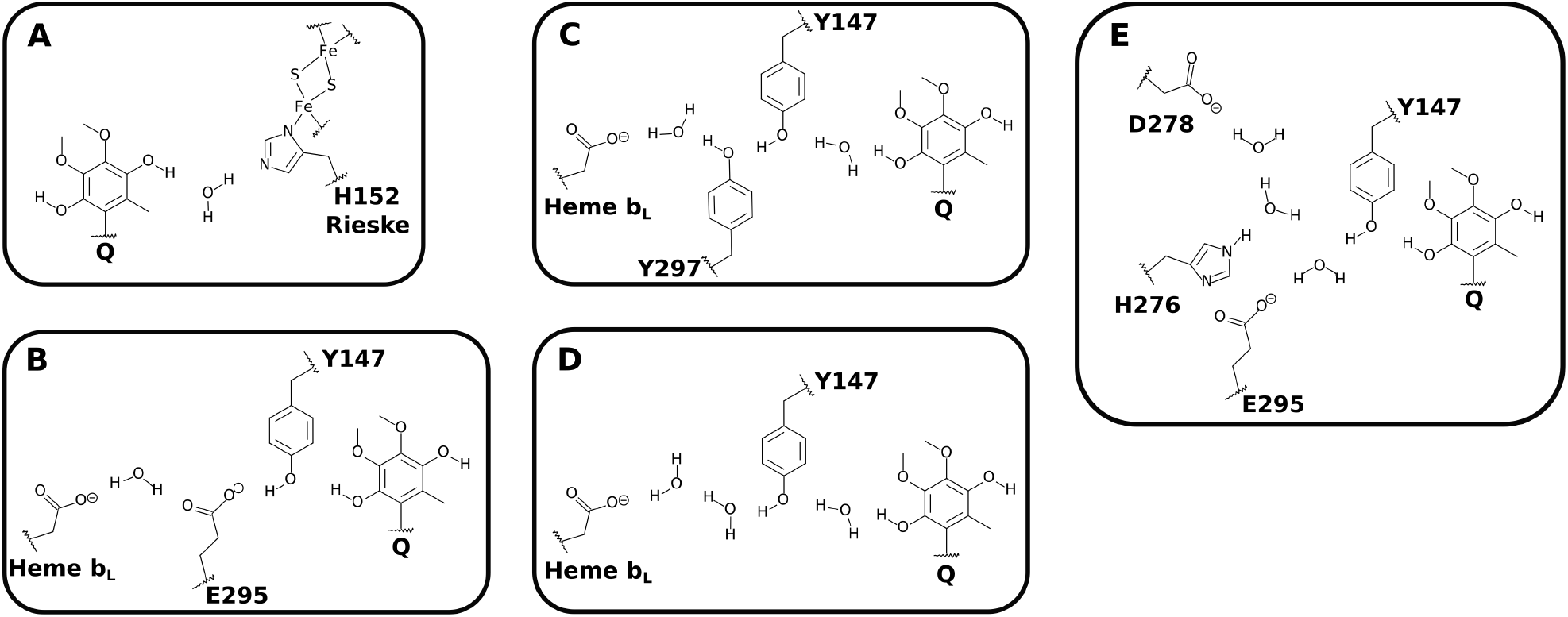
Proton-wires proposed to transfer the two chemical H^+^ generated by oxidation of the quinol substrate to proton release groups. **A** shows the pathway for proton transfer from Q_*O*4_ to H152 in the Rieske protein. **B**-**D** show pathways from Q_*O*1_ to heme PRA_*bL*_. **E** shows the pathway from Q_*O*1_ to D278 in cyt *b*.

Contacts between heme PRA_*bL*_ and residues of the YEY group can only be established via water bridges. On the other hand, contacts between Q_*O*4_ and H152, Q_*O*1_ and Y147, Y147 and E295, and between H276 and D278 may be established directly or via water bridges (Fig. 5 and Fig. 9). To establish the preferred proton-wire or the role of the multiple ones found here, their exact composition and the number of participating water molecules necessary for efficient proton conduction requires further simulations of the proton transfer reaction during Q oxidation, such as using hybrid QM/MM potentials,^36,65,66^ and are outside the scope of this study.

In the Q_*o*_ site, there is an interplay of direct H-bonds or contacts mediated by water molecules in bridges (H-bonded in series) due to the dynamic network of H-bonds established. We refrained from proposing wires with connections containing more than two water molecules in a bridge. It is expected that the stability of long wires will decrease with an increasing number of mobile participants, such as bridge waters, due to the entropic penalty of having all groups aligned simultaneously.^31,67^ Yet, the presence of multiple wires may reduce this entropic penalty and avoid disruption of function in point mutants of participating residues.

Site-directed mutagenesis at the Q_*o*_ site has been widely studied in experiments aimed at determining residues involved in proton transport.^4,68^ However, their interpretation must consider that point mutations may perturb local conformations (or even global protein stability and assembly) and lead to the indirect deactivation of proton transfer pathways without necessarily knocking out a residue that chemically reacts. Also, proton transfer is often fast, may not be rate-limiting, and may operate through various proton-wires. Thus, multiple mutations may have to be combined to significantly slow or alter measured kinetics and organism growth properties.^30^ These multiple mutations may also exacerbate the indirect perturbation effect.

Yet, we should mention a few mutational studies in line with the discussion presented here. Exchange of E295 affects overall *bc*_1_ catalytic activity but does not completely abolish it.^29,69–71^ Mutation of Y147 is more drastic, and the *bc*_1_ activity is severely affected, in some cases making electron transfer from quinol to heme *b*_*L*_ impossible.^28,68,72^

One of the open questions about the mechanism of Q oxidation in the Q_*o*_ site is the relevance of a semiquinone (radical) intermediary for the bifurcated electron transfer. Although this radical has been experimentally reported,^22,73^ the evidence has been debated, particularly in the context of wild-type *bc*_1_ under normal operation of the Q-cycle.^4^ The present results show that pathways for proton transfer from Q_*O*4_ and Q_*O*1_ reform on the ns time scale due to side chain torsions, Q-head flexibility, and local hydration in the Q_*o*_ site. We suggest this time scale is a lower bound to the lifetime of the semiquinone intermediary. Two-electron oxidation of the quinol substrate without proton transfer (or generation of a QH^2+^ transient species) is highly unlikely, so at least one concerted proton transfer^26^ should take place during the oxidation process, requiring the formation of a proton-wire on the ns time scale.

### Experimental structures confirm that Q_*o*_ side chains are flexible and highly hydrated

We collected 79 experimental structures of the Q_*o*_ site previously deposited in the PDB to analyze the distribution of side chain conformers and local hydration (see details in “Analysis of experimental structures” in Methods and “Details of *bc*_1_ experimental structures analyzed here” in the SI). Table 1 and Fig. 10 show the five principal torsional modes of YEY side chains found experimentally. The most frequent modes (A and B) have Y147, E295, and Y297 *χ*_1_ in the (*t*, −*g, g*) conformer and differ only in the E295 *χ*_2_ torsion, with this side chain pointing towards the Q substrate (B) or in the opposite direction (A). Other conformers are observed for YEY side chains, particularly for Y147, which may assume all three conformers, in agreement with our simulation results on the flexibility of these side chains.

**Table 1:**
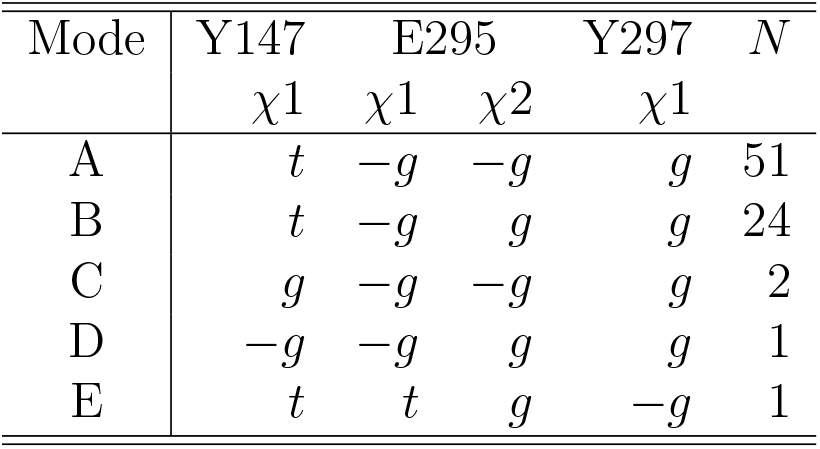
Experimental torsional modes for YEY residues shown in Fig. 10 and found in *N* structures of the experimental set.

**Figure 10:**
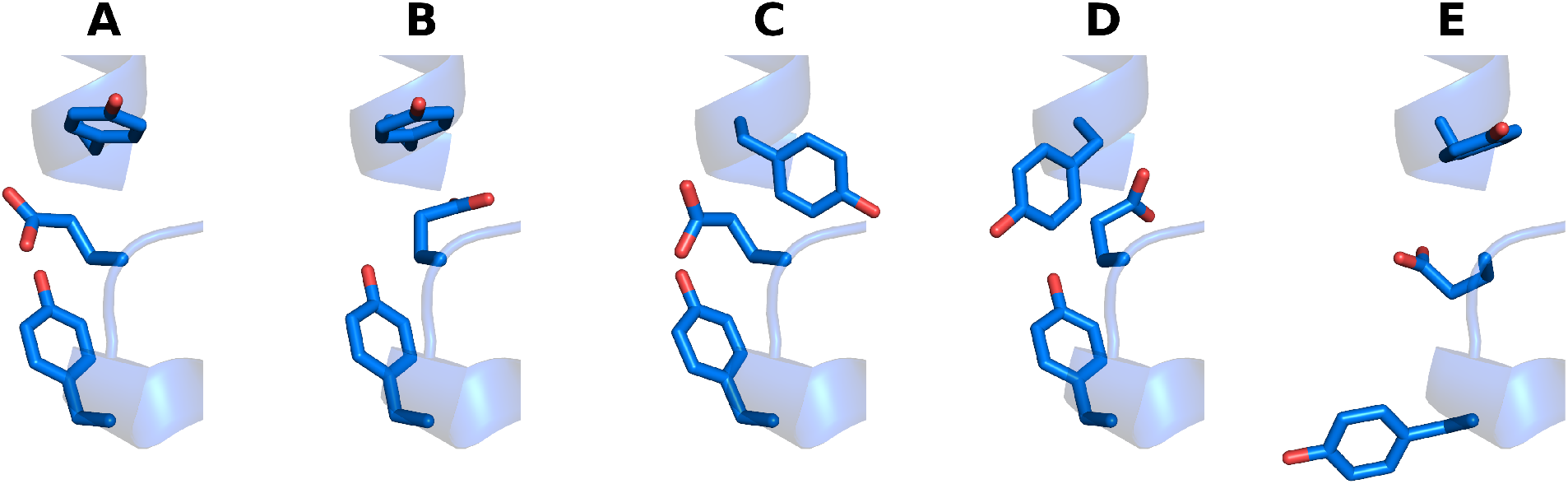
Experimental conformations for Y147, E295 and Y297 (YEY) residues in the Q_*o*_ site found among 79 structures of the *Q*_*o*_ site deposited in the PDB. Modes are labeled **A**-**E** accordingly to Table 1.

**Figure 11:**
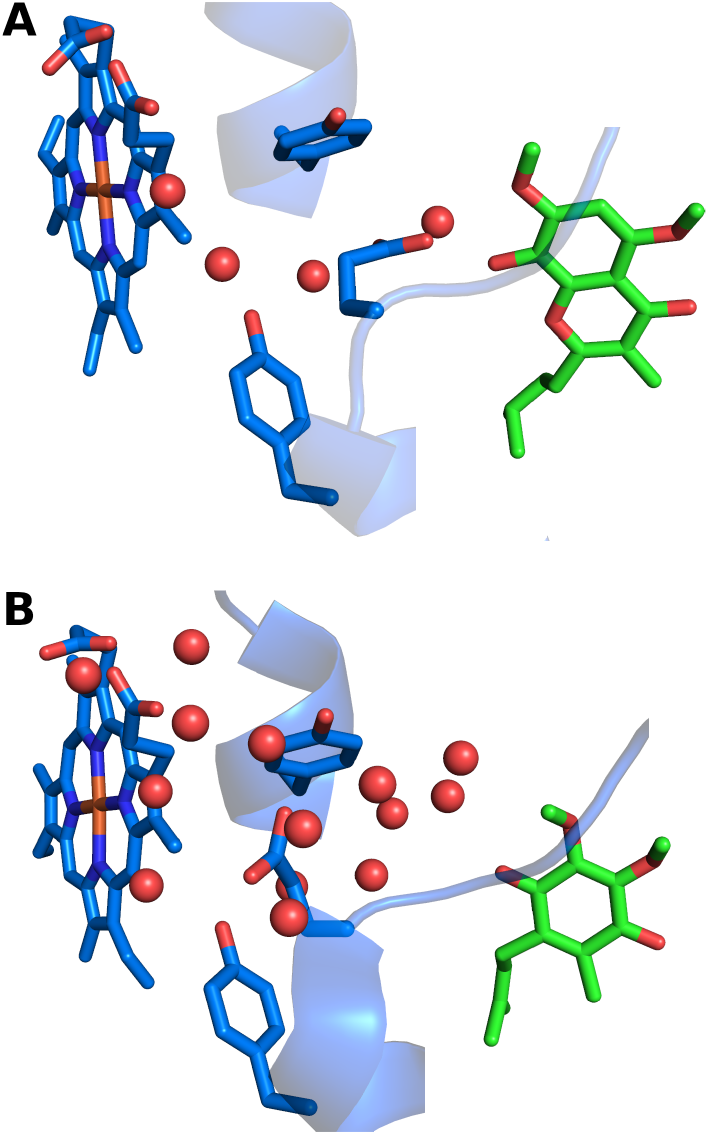
Water molecules found in experimental structures of the Q_*o*_ site. **A** shows PDB 1sqx^74^ with water between Y297 and heme PRA_*bL*_ and between the bound stigmatellin inhibitor and E295. **B** shows PDB 8bel^16^ also with water between Y147, E295 and distal bound Q-head substrate, in line with our MD simulations.

It is remarkable that the relative population among binding modes found experimentally is in line with the relative free energies estimated from simulations. For instance, the minima found in the 2D free energy profile for combined Y147 and Y297 torsion (Fig. 3D) directly correspond with experimental modes (Table 1) and conformers shown in Fig. 3G,H,I map respectively to modes (A,B), D, and E in Fig. 10. Relative conformer populations of 1-2%, observed for modes C-E (Table 1), correspond to differences in free energy of ∼10 kJ/mol between their relative stability, as estimated here (Fig. 3D). The torsion profile for *χ*_1_ in E295 (Fig. 3F) shows the *g* conformer is ∼20 kJ/mol less stable, and this form is not observed in the experimental set.

Given the high variability in the residues found at cyt *b* positions 276 and 278 (Fig. 2), H276 and D278 are both present in only half (40) of the 79 experimental Q_*o*_ site structures analyzed. D278 *χ*_1_ assumes conformer −*g* in 35 structures. H276 *χ*_1_ is more flexible and assumes either −*g* (19 structures) or *t* (21 structures) forms. In the H276 −*g* torsion, the distance between H276_*N*(*δ,ϵ*)_ and D276_*Cγ*_ is higher than 0.7 nm, and this conformer should be less relevant for the proposed proton-wire (Fig. 9E). The distance is reduced to 0.5 nm in the *g* H276 conformer, shown in simulations to be energetically stable (Fig. 3E).

Among the experimental structures with resolved water in the Q_*o*_ site, up to 5 water molecules were found within a 0.5 nm radius of heme PRA_*bL*_, 4 near Y147, and 3 near Y297, indicating hydration in the Q_*o*_ site region and the possibility of water-mediated contacts and proton transfers. Water-bridged contacts are revealed in various structures.

For instance, an X-ray structure bound to stigmatellin (Fig. 11A) shows water molecules between heme *b*_*L*_, Y297, and E295. A recent structure of the supercomplex I + III_2_ of *Arabidopsis thaliana* in high resolution shows extensive hydration of the Q_*o*_ site^16^ (Fig. 11B), and reveals a pocket of water molecules near PRA_*bL*_, drawing attention to the role of Y297 and nearby water molecules in mediating proton transfer.

### Conclusions

Molecular dynamics simulations of cytochrome *bc*_1_ provide insights into the conformational landscape of the Q_*o*_ site and its binding to the Q substrate in quinol form. Conserved side chains (YEY group of Y147, E295, and Y297) populate multiple conformers, resulting in three binding modes for the Q-head. The two modes proximal to heme *b*_*L*_ may transfer protons upon Q oxidation via direct and water-bridged interactions between Q_*O*1_ and Y147, and between Q_*O*4_ and H152, suggesting these residues are initial proton acceptors. The distal binding mode is characterized by a lack of contact between Q_*O*1_ and Y147, and is in line with the Q binding mode observed in experimental structures, likely representing a stable pre-reactive state that is more readily captured in cryo-EM studies.

Analysis of a collection of experimental structures confirms the flexibility of YEY group, particularly for the Y147 and Y297 side chains. This flexibility has not received much attention before. Simulations also indicate that H276 and E295 form weak interactions within the Q_*o*_ site and are not initial proton acceptors. Instead, H276 may only engage in indirect proton transfer from Q through Y147 and E295.

MD simulations also reveal that the Q_*o*_ site is highly hydrated, with a network of both direct and water-mediated contacts. Key residues such as Y147 and H152 also interact with the Q substrate through these water-bridged connections. PRA_*bL*_ only interact with YEY residues via bridged contacts. Thus, multiple proton-conducting wires are established for efficient transfer of protons in a Grotthuss mechanism, highlighting the critical role of hydration in the Q_*o*_ site function. This is again supported by experimental structures, particularly recent cryo-EM studies.

Our results agree with the proposed roles for H152 in the Rieske protein for Q binding in the Q_*o*_ site and transport of one of the chemical protons released from Q oxidation.^2,4,20^ From H152, the excess proton could be passed to bulk water during the characterized movement of the Rieske protein, that approximates the FeS cluster and heme *c*_1_ to continue the electron transfer chain.^4^

An important contribution of this study concerns the groups and proton-wire involved in transport of the secondary proton from Q oxidation. Our results strongly support Y147 as the initial proton acceptor from Q_*O*4_, and we propose for the first time that heme *b*_*L*_ propionate-A is the final protein acceptor before the excess proton is released to the bulk water. An alternative proposal^30^ suggesting a proton-wire composed of H276 and D278 seems less likely due to lack of their conservation and the complex, extended pathway (Fig. 9E) required.

Multiple proton-wires and a network of H-bonds have been proposed to transport protons in other proteins^75,76^ including respiratory complex IV (CcO) which also show proton-wires composed by propionate groups from a redox-active heme.^33–35^ Thus, it is not really surprising that cytochrome *bc*_1_ employs similar components and interactions for the molecular mechanism of the Q-cycle in the Q_*o*_ site as suggested here by MD simulations. The correlation between our detailed results and experimental data strengthens our understanding of the molecular mechanisms governing proton transfer in cytochrome *bc*_1_, integrating structural flexibility with functional hydration.

In closing, it is notable that other enzymes involved in Q redox reactions appear to use similar residues and strategies for transferring chemical protons to the substrate Q. Respiratory complex I (NADH:ubiquinone oxidoreductase)^61,62^ and complex II (succinate dehydrogenase)^77^ also contain His and Tyr side chains that hydrogen bond to Q molecules, either directly or via water bridges, within their active sites where Q redox reactions occur. This similarity warrants further investigation to elucidate a general enzymatic mechanism for Q redox chemistry.

## Associated Content

### Data Availability Statement

All data related to this study is available from the corresponding author upon reasonable request by email.

## Supporting Information

Details of *bc*_1_ experimental structures analyzed here, root mean square deviations (RMSD) and fluctuations (RMSF) from MD simulations, convergence of metadynamics simulations and free energy profiles, trajectories of contacts by heme PRA_*bL*_ and by H276, and by additional contacts bridged by water, and supporting references for experimental structures analyzed.

## Acknowledgement

Funding from Fundação de Amparo à Pesquisa do Estado de São Paulo (FAPESP, scholarship 2019/05531-0 to S.R.G.C and grants 2019/21856-7 and 2023/00934-5 to G.M.A.) and computational resources from the SDumont cluster in the LNCC (MCTI) are gratefully acknowledged. The authors declare that they have no conflicts of interest with the contents of this article.

