## Supporting Information file for "Flexibility and hydration of the Q_*o*_ site determine multiple pathways for proton transfer in cytochrome *bc*_1_"

**Contents:**

- Supplementary Text: Details of $bc_1$ experimental structures analyzed here
- Figure S1: Root mean square deviations (RMSD) from MD simulations
- Figure S2: Root mean square fluctuations (RMSF) from MD simulations
- Figure S3: Convergence of metadynamics simulations and free energy profiles
- Figure S4: Trajectories of contacts by heme $b_L$ A-propionate and by H276
- Figure S5: Trajectories of additional contacts bridged by water
- Supporting References

### Details of $bc_1$ experimental structures analyzed here

In order to probe the conformational distribution of residues in the $Q_o$ site using high-resolution experimental data, 52 entries of cytochrome $bc_1$ and its $b_6f$ analogue were retrieved from the PDB with a resolution better than 4.0Å. An initial set was obtained in 2021 when this analysis was first carried out. Various cryo-EM structures containing the $bc_1$ dimer were obtained more recently and all structures containing a bound Q substrate were added to this analysis resulting in 79 models of the $Q_o$ site.

The $Q_o$ experimental structures were classified into five conformational modes based on the side chains of Y147, E295, and Y297 (Fig. 10 and Table 1). Modes A and B together consist of 75 $Q_o$ site models from various organisms. In Mode A, there are 51 structures predominantly from *Bos taurus* (12), *Saccharomyces cerevisiae* (10), *Arabidopsis thaliana* (6), *Mus musculus* (4), *Vigna radiata* (4), and *Sus scrofa* (4). In Mode B, 24 structures come primarily from *R. sphaeroides* (10), *Gallus gallus* (4), and *Saccharomyces cerevisiae* (4). The majority of these structures belong to the cytochrome $bc_1$ complex, with 50 structures alone, 22 as part of supercomplexes containing $bc_1$, and 3 of cytochrome $b_6f$. 55 models do not have resolved water molecules. Cryo-EM was used to determine 38 of these structures, while X-ray crystallography was used for 37. Regarding ligand presence at the $Q_o$ site, 27 structures in Mode A have various ligands, while 21 structures in Mode B specifically have stigmatellin-A as a ligand, which is consistent with its role in inducing conformational changes at E295.[4]

In the mode C, there are only two structures from *Bos taurus* determined by X-ray crystallography without resolved water molecules. One structure contains the ligand *6-hydroxy-5-undecyl-1,3-benzothioazole-4,7-dione* (UHD). Mode D consists of only one structure of the cytochrome $b_6f$ from *Chlamydomonas reinhardtii* determined by X-ray crystallography and does not contain resolved water molecules. The ligand at the $Q_o$ site is *8-hydroxy-5,7-dimethoxy-3-methyl-2-tridecyl-4H-chromen-4-one* (TDS). Mode E also includes only one structure of the cytochrome $b_6f$ complex obtained from *Mastigocladus laminosus.* This structure does not contain resolved water molecules or ligand at the $Q_o$ site.

In the 79 models of the $Q_o$ site, 13 are occupied by Q (PDB entries: 1ntz, 6q9e, 7rja, 8asi, 8bel, 8bpx, 8bq5, 8bq6, 8e7s[13–19,78]), with one site occupied by ubiquinone-2 ($UQ_2$), four by $UQ_5$, two by $UQ_6$, four by $UQ_7$ and two by $U_{10}$. All sites occupied by Q are part of mode A.

The following PDB entries were analyzed: 1kb9, 1kyo, 1l0n, 1ntk, 1ntz, 1p84, 1q90, 1sqb, 1sqp, 1sqq, 1sqv, 1sqx, 2d2c, 2e76, 2fyn, 2fyu, 2ibz, 2qjk, 2qjp, 2qjy, 2ybb, 2yiu, 3h1i, 3h1j, 3l72, 4h13, 4pd4, 5kli, 5klv, 5nmi, 6fo6, 6hu9, 6kls, 6nhg, 6q9e, 6rqf, 6t0b, 7jrg, 7jrp, 7o37, 7o3c, 7o3e, 7o3h, 7rja, 7tlj, 8asi, 8bel, 8bpx, 8bq5, 8bq6, 8e7s, 8uge, 8ugf, 8ugg.[9,14–17,19,58,74,78–100]

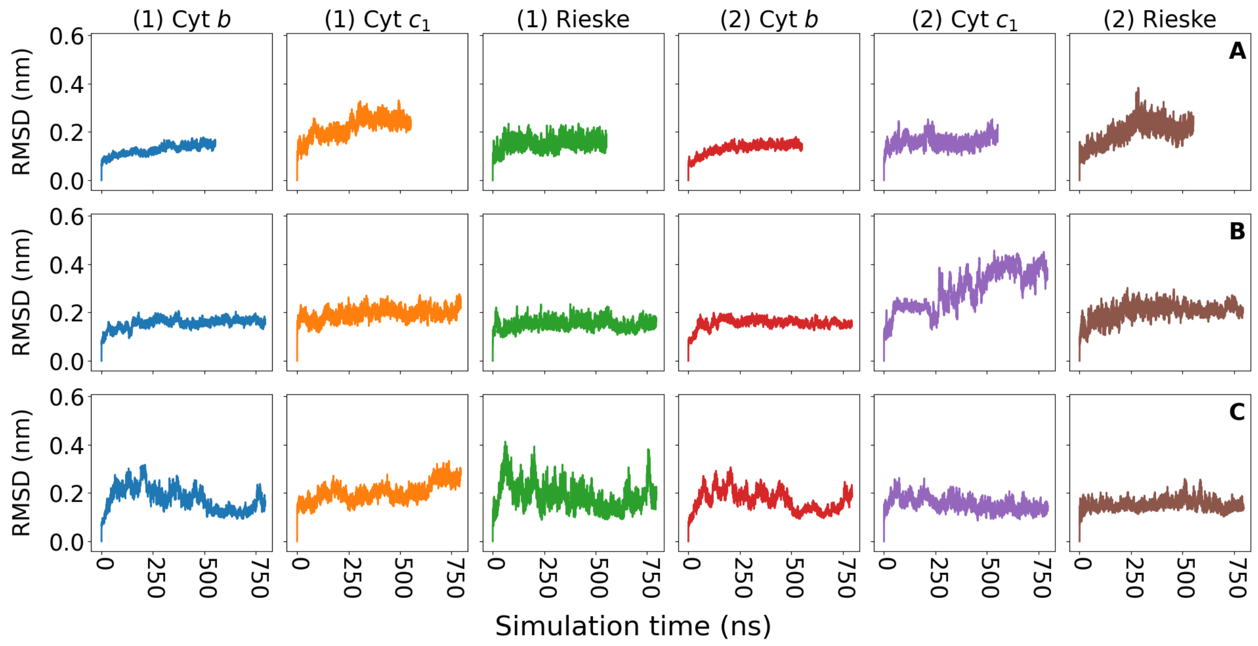


Figure S1: Root mean square deviations (RMSD) of C$\alpha$ atoms obtained from MD simulations of the cytochrome $bc_1$ dimer, for each protein subunit named on top of each column. **A** shows the 550 ns canonical MD. **B** has metadynamics activated for the YEY group of the $Q_o$ site in chain (1)-Cyt *b*. **C** has metadynamics activated for the YEY group in chain (2)-Cyt *b*. All simulations remain stable. The transition observed in panel line B for (2)-Cyt $c_1$ around 300-500 ns corresponds to residues 185 to 220 (high RMSF in Fig. S2B), a flexible region exposed to solvent and without stable secondary structure.

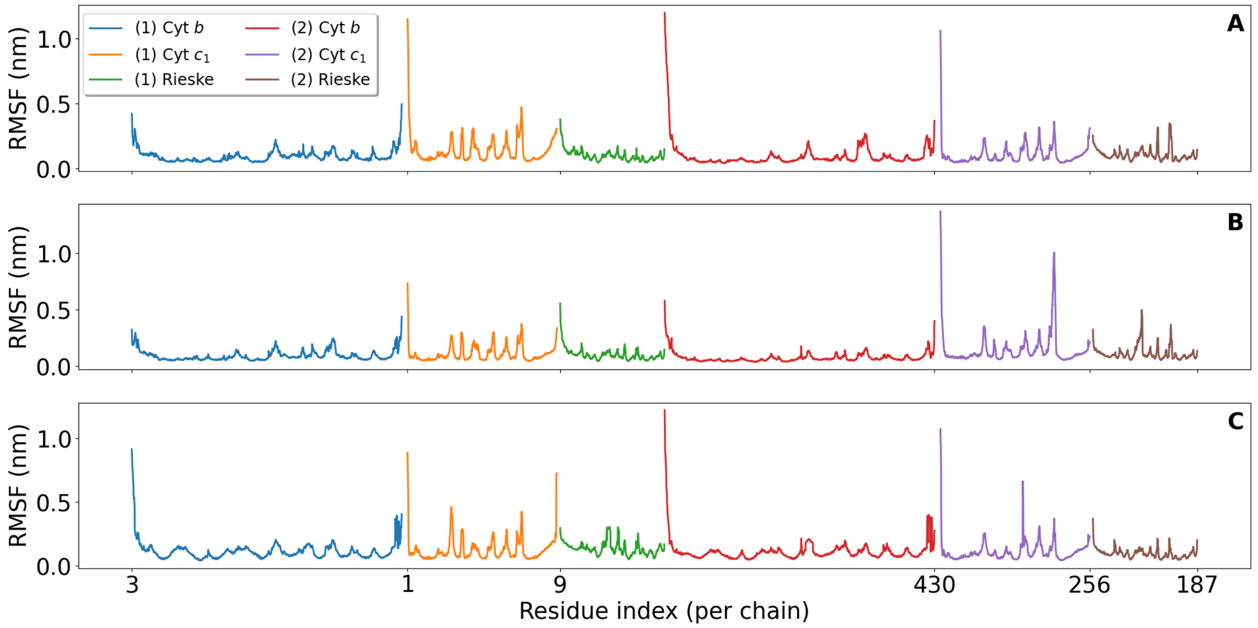


Figure S2: Root mean square fluctuations (RMSF) of backbone heavy-atoms obtained from MD simulations of the cytochrome $bc_1$ dimer. **A** shows the 550 ns canonical MD. **B** has metadynamics activated for $Q_o$ site in chain (1)-Cyt *b*. **C** has metadynamics activated for $Q_o$ site in chain (2)-Cyt *b*. Initial and final residue indexes are shown for subunit sets (1) and (2), respectively.

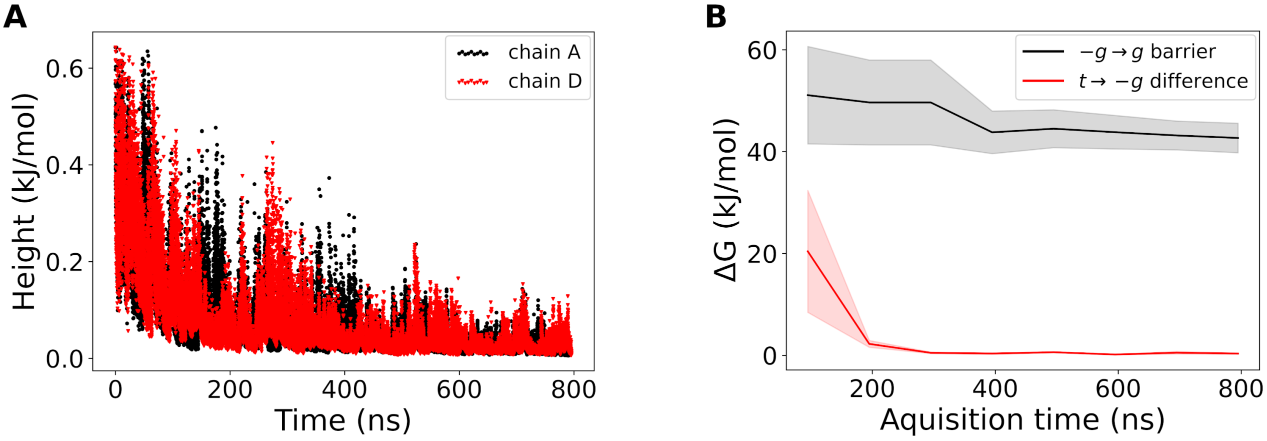


Figure S3: Convergence of the metadynamics simulations and derived free energy profiles. **A** shows the time evolution of the gaussian height in the two well-tempered metadynamics simulations performed (Fig. 3). The height decreases and levels off around 400 ns, suggesting that a smoother energy surface is visited and enhanced sampling of the YEY side chain dihedrals is achieved. **B** shows the convergence with simulation time of three free energy differences between Y147$_{\chi 1}$ conformations indicated in the legend. The average between the two simulations is shown with colored shadows giving the standard error. Variations in free energies are smaller than this statistical error after 400ns, indicating convergence. A similar behavior is observed in free energy profiles for the other boosted YEY dihedrals.

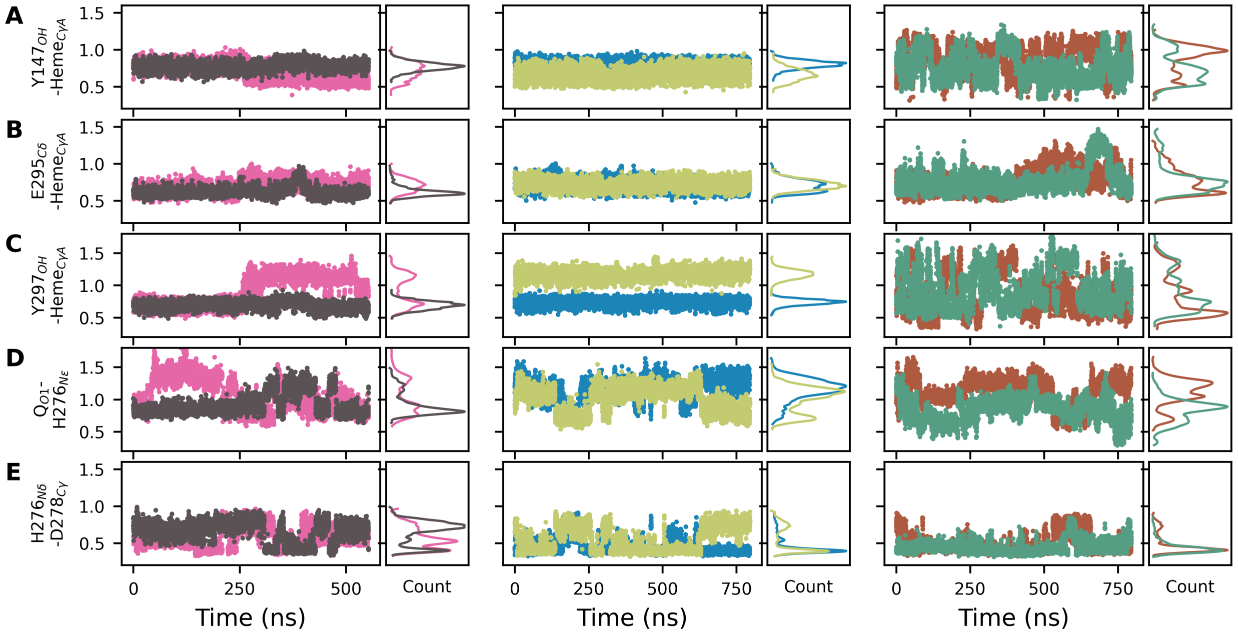


Figure S4: Atom-pair distances involving heme PRA$_{bL}$ in **A**-**C** and H276 in **D** and **F**. Pairs are given in Y-axis labels. Panel columns and colors relate to the six MD simulations of the Q$_o$ site as described in Fig. 4. Count shows a histogram of the respective distance.

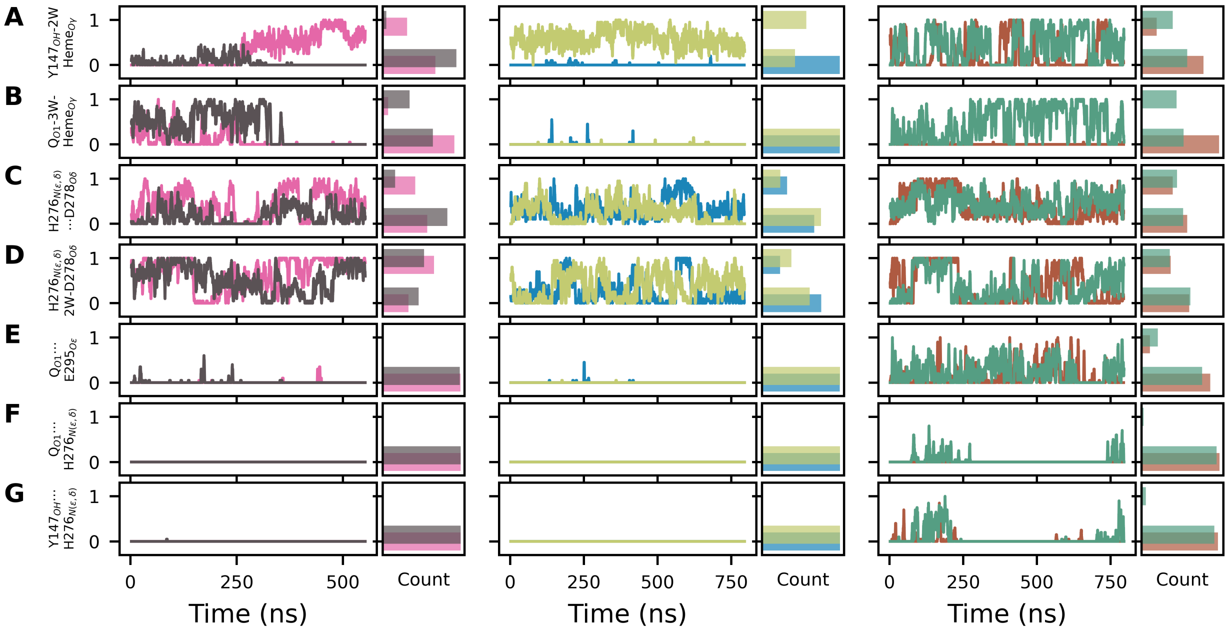


Figure S5: Additional contacts bridged by water, as labeled in Y-axis. Dots ($\cdots$) in labels correspond to one bridge water molecule. Bridges with two or three waters are labeled 2W or 3W, respectively. One or zero correspond to the contact formed or not, respectively. A moving-average with a 1 ns window is plotted. Count shows a histogram of contact formation. Panel columns and colors relate to the six MD simulations of the $Q_o$ site as described in Fig. 4.
